# Using Recognition Testing to Support Semantic Learning in Developmental Amnesia

**DOI:** 10.1101/2023.03.13.532399

**Authors:** Rachael Elward, Jennifer Limond, Loïc Chareyron, Janice Ethapemi, Faraneh Vargha-Khadem

## Abstract

Patients with developmental amnesia (DA) have suffered hippocampal damage in infancy and subsequently shown poor episodic memory, but good semantic memory. It is not clear how patients with DA learn semantic information in the presence of episodic amnesia. However, patients with DA show well-developed recognition memory and these recognition abilities may support semantic learning. We present data from three experiments (two previously described in Elward & Vargha-Khadem, 2018). The first experiment showed that recall tests did not facilitate semantic learning. Patients with DA recalled only 35% of the learned information (controls recalled 80%). The second experiment indicated that multiple-choice recognition tests may facilitate learning. Patients with DA recalled 85% of the learned information. In experiment three, a patient with DA (aged 8 years) took part in a repeated-measures test so that recall learning and recognition learning could be directly compared. The results showed a clear benefit of recognition learning compared to recall learning (76% v. 35%). This finding indicates that young people with extensive hippocampal damage indeed utilise their recognition memory to support the integration of new information into their semantic system. This has important implications for the support of school-aged children with episodic memory difficulties.

## Introduction

Children with developmental amnesia (DA) have suffered a hippocampal injury in early life. Although there are a range of aetiologies, a common cause is a traumatic birth and asphyxia causing a hypoxic-ischemic injury in the neonatal period (Gadian et al., 2000; Vargha-Khadem et al., 1997). After recovery from the initial trauma, early development appears normal. Children with DA meet the expected developmental milestones in infancy and early childhood for motor, language and cognitive development. However, difficulties with memory emerge during the preschool years. Parents and caregivers notice that the child does not remember events that they are expected to remember, including birthday parties and memorable occasions. They cannot provide a reliable account of the day’s activities, remember conversations, retell stories etc. Children with DA often lose their belongings and can get lost in familiar surroundings. These memory difficulties interfere with daily living to the extent that children and adolescents with DA cannot gain independence commensurate with their age and aspirations. They depend heavily on support from family members and teachers.

A fascinating feature of developmental amnesia is that semantic memory (that is, memory for factual knowledge) is spared (Elward & Vargha-Khadem, 2018). Children with DA have age-appropriate world knowledge that they can express through their well-developed speech and language abilities (Gadian et al., 2000; Vicari et al., 2007). In fact, one report indicates that semantic knowledge in adults with a history of DA can be superior to that of matched controls (Jonin et al., 2018). Semantic memory is a strength that can be used to compensate for some aspects of the developmental amnesic syndrome. For example, if the child cannot recall a specific event, they may use their semantic memory to guess at an appropriate response (Brandt et al., 2006).

Despite this strength, children with DA struggle to keep up with peers in education. In laboratory studies, patients with DA take longer to learn semantic information than controls (Baddeley et al., 2001; Gardiner et al., 2008). It’s likely that typically-developing children use their episodic memory (viz. autonoetic memory of their life experiences) to recall previous lessons and learning episodes and use this to support newly-learned information before it is consolidated into the semantic system. Patients with DA do not have access to episodic memory and so are at a considerable disadvantage when learning new information. As a result, young people with DA find school a struggle and fall behind their peers. In secondary education and beyond they tend to take up vocational skills training instead of formal education, even if this is not in-line with their interests and cognitive abilities. Anecdotally, young people with DA fail to thrive in education and do not find fulfilling employment in adulthood.

This patient group has inspired scientific research; however, the academic community has not been able to develop rehabilitative techniques that can support children with DA to reach their potential in education and employment. Some techniques that originally seemed promising (i.e. the fast-mapping technique) have failed to support learning in DA (Elward et al., 2019). From the research that has been conducted, three conclusions can be drawn about the optimum conditions for learning in DA. The first of these findings is that patients with DA will benefit from repeated presentations of the study material. Baddeley at al., (2001) demonstrated that when patient Jon was shown four presentations of previously unfamiliar newsreel footage, he performed as well as controls in an immediate recall test, however, underperformed relative to controls in an overnight recall test. Second, patients seem to benefit from consolidation over time, or rather, healthy controls show more forgetting over time than patients, and therefore testing memory after a delay of several days or weeks will show smaller group differences than testing on the same day (Elward & Vargha-Khadem, 2018; Gardiner et al., 2008). Lastly, patients with DA perform well on recognition memory tests. That is, when given a multiple-choice test in which the patient needs to only recognise the familiar option from a list of alternatives, then patients with DA perform as well as their peers.

However, when given a free recall test, in which the patient must bring back to mind the relevant information from memory, the patients with DA markedly underperforms (Baddeley et al., 2001; Gardiner et al., 2006). This last finding may be crucial for supporting learning in patients with DA. It demonstrates that patients with DA show some evidence of semantic learning but that the memory is weak and accessible to recognition only (Elward & Vargha-Khadem, 2018, Vargha-Khadem et al. 2006). The key to promoting semantic learning in DA may be to consider how this nascent memory may be strengthened so that patients can retrieve the information that they have learned through recall.

In our review, we reported unpublished data from Limond and colleagues suggesting that testing through recognition can indeed support semantic retrieval in DA (Elward & Vargha-Khadem, 2018). We compared data from two previously unpublished studies completed by Limond and colleagues. In both studies, patients with DA and controls were asked to learn novel semantic information in the form of short narratives. The information was presented six times, and after each presentation, the patient underwent a memory test. The key difference between the two studies was the type of memory test that was used to support learning: A recall test or a recognition test.

In the first experiment participants completed a recall test after each presentation of the semantic information (see Figure 1, Recall Learning). In this condition, the patients underperformed relative to controls on all the learning trials and the delayed recall test (see Figure 1: cued recall test, highlighted with a box).

**Figure 1:**
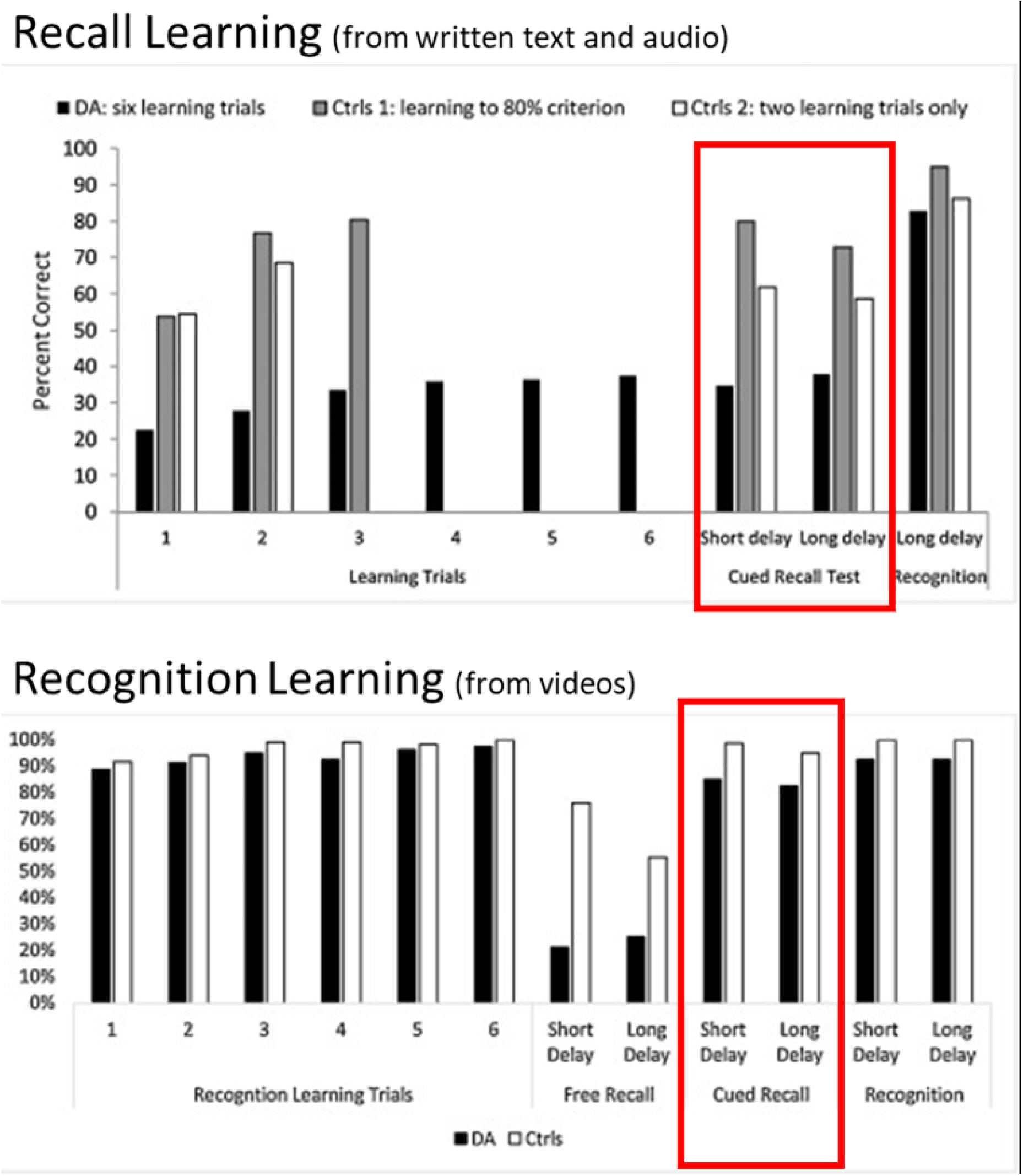
Reproduced from Elward & Vargha-Khadem (2018). Data from two studies that designed to investigate semantic learning in DA. In Recall Learning (top panel) participants were asked to complete six recall tests during learning. The cued recall test (highlighted in with a box) indicates that participants did not show good learning with this method. In recognition learning (bottom panel) participants were asked to complete recogntion tests during learning. In this case, performance on the cued recall test was similar to controls.

The second experiment (see Figure 1, Recognition learning) was conducted several years later. This time the participants learned semantic information from videos instead of text, but the same semantic information was used. After each presentation of the video, participants completed a recognition test (i.e., a multiple-choice test). As expected from the typical profile of DA, the patients with DA performed well on these recognition tests during the learning trials. The true test of learning occurred one week later in a delayed recall test (see Figure 1: cued recall test, highlighted with a box). In the recognition learning experiment, patients with DA were able to recall the semantic information at a level comparable to controls. In fact, patients recalled twice as much information in this condition as the previous version of the experiment. We interpret that having repeated opportunities to recognise the correct information during the learning strengthened the semantic memory and facilitated recall.

This finding is encouraging but must be interpreted with caution. Some of the same patients participated in both the recall learning experiment and the recognition learning test several years later. Therefore, it is possible that participants had consolidated the information from the recall learning experiment in the intervening years and if so, this would explain their improved performance on the later test.

Furthermore, the first experiment used written materials and audio recordings, so learning the semantic information relied only on auditory/phonological processing, but the second experiment used videos which contained visual images too. It’s not possible to determine if recognition testing is driving the new semantic learning, or if the addition of visual stimuli is key. In addition, the DA patients were adults at the time of the recognition learning experiment and so it’s not clear that this method would support learning in school age children where it could have the biggest impact on education and life success.

In order to test the hypothesis that recognition-based learning will facilitate semantic learning in children with DA, here we report quai-experimental (pre-post) design in which a child with DA (aged 8) encounters new semantic information (presented via videos) for the first time. Across three weeks of testing, two videos were presented in a recognition learning condition where learning is supported with multiple choice tests, and two were presented in the recall learning condition (where learning is supported with open ended questions). The outcome measure is cued recall performance after a delay of one week.

## Methods

### Participants

#### Case Presentation

Patient H, an eight-year old boy, was referred by his General Practitioner to a Consultant Neuropsychologist (FVK) at Great Ormond Street Hospital NHS Trust due to concerns about his “excessive forgetfulness”.

During the clinical interview, Patient H’s mother reported that her son had a complicated birth involving meconium aspiration and asphyxia. Shortly after birth he was transferred to intensive care where he was ventilated for three days, and then remained in hospital for two weeks. Following this stormy period, H’s condition improved and at discharge he showed no neurological symptoms. Patient H met his developmental milestones as expected. His mother described him as an easy baby who did not raise any concerns until the age of three when she noticed his difficulties remembering daily events. These concerns became more prominent when Patient H started school and had difficulty settling in; he was distressed, could not remember where his mother was and asked to go home. In school, he could not follow the teacher’s instructions and soon started to fall behind his peers. Teachers shared his mother’s concerns about his memory. Apart from the memory difficulties, H’s mother does not report any other cognitive difficulties or behavioural problems. She describes her son as a healthy, bright, and sociable boy who has good general knowledge and language skills. She remarked that some aspects of Patient H’s memory seem to be working well as he had learned the names of many Pokémon characters despite his amnesia.

### Brain Volume Measurements

A T1-weighted MRI scan (flip angle of 8°, field of view = 25.6 cm, repetition time = 2300 msec, and 1 mm isotropic voxels) was acquired on a Siemens 3-T Prisma scanner equipped with a 32-channel receiver head coil at Great Ormond Street Hospital for Children NHS Foundation Trust, London, UK. The clinical neuroradiology report indicated that the hippocampi were small and the mamillary bodies were visible, but atrophied. No other abnormalities were identified.

Manual segmentation of the MRI acquisitions was used to estimate the volume of the hippocampus as a whole and several subregions, namely the uncus, CA-DG (including CA fields and dentate gyrus) and subicular complex (including subiculum, presubiculum and parasubiculum). These subregions are plotted alongside those obtained from the group of controls shown in Fig 2.

**Figure 2:**
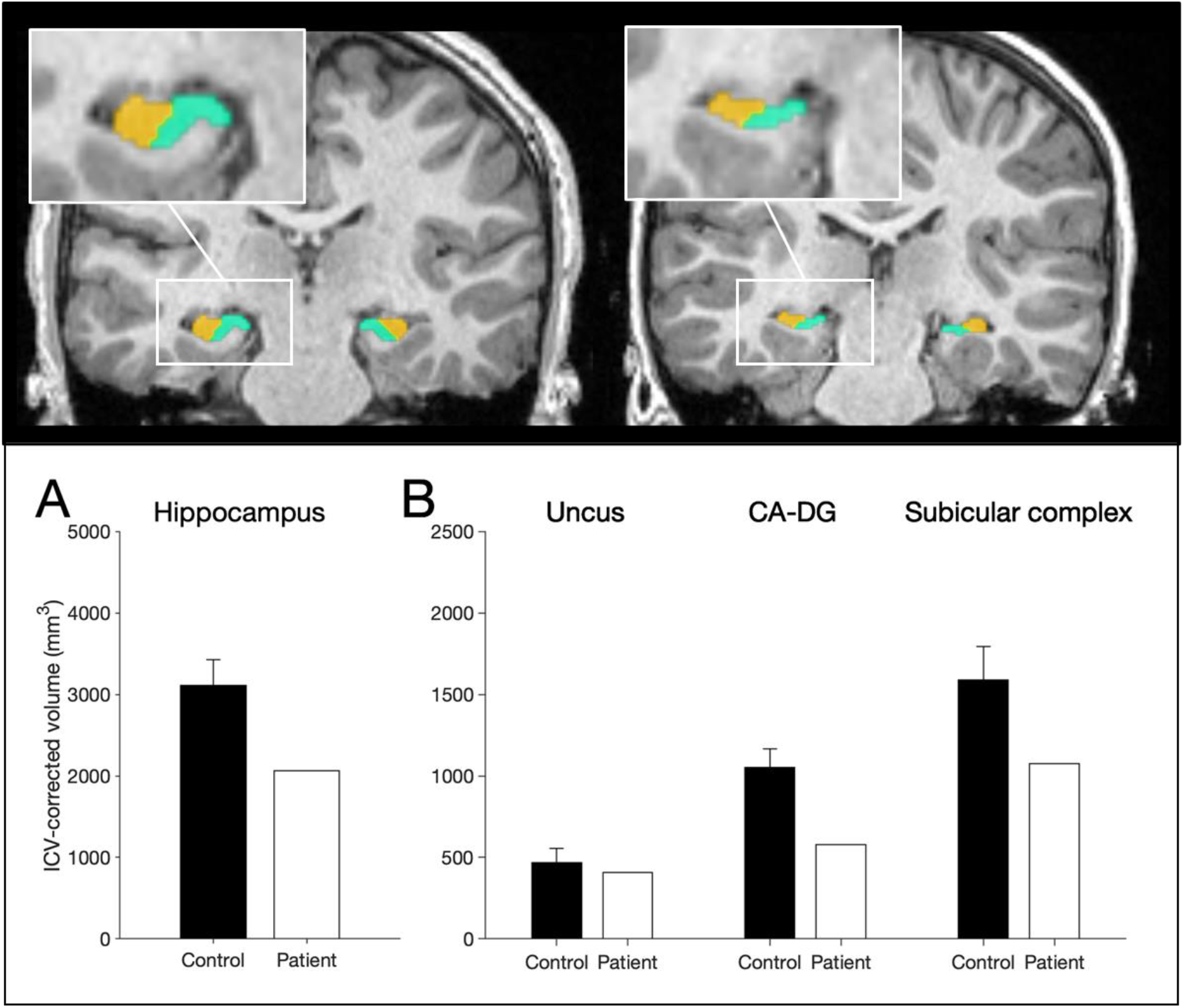
Manual segmentation of the hippocampus. Top: 3T-MRI scan of a 14-year-old male control participant (left) alongside Patient H (right) showing the reduced volume of the subicular complex (in green) and CA-DG region in (orange). Bottom. ICV-corrected hippocampal volumes for patient H compared with 32 healthy controls (8y – 38y; 16male) indicating a 34% volume atrophy of the hippocampus.

### Typically Developing Controls

Five new control children (1 male) between ages of 8–10 years were recruited through the local community at UCL and LSBU. The inclusion criteria were that children must be aged between 7–10 years of age, the product of normal birth, generally healthy with no medical conditions and no learning difficulties.

Control children participated in a short neuropsychological assessment consisting of the Weschler Abbreviated Scale of Intelligence (WASI-II) and the Children’s Memory Scale. The five controls performed within the normal range on all subtests of each test. Full scale IQ ranged from 112 to 134 (mean 124).

The research was overseen by the Research and Development Department of Great Ormond Street Hospital for Children NHS Foundation Trust, and the UCL Great Ormond Street Institute of Child Health, London, UK. The project was approved by the Hampstead NHS Research Ethics Committee and was also approved by the Ethics committee of School of Applied Sciences, London South Bank University. Testing occurred in a laboratory space at the Wolfson Centre laboratories at the UCL Great Ormond Street Institute of Child Health or in the Child Development Laboratory at London South Bank University. Each child was accompanied to the testing sessions by a parent who provided informed consent and waited in a nearby waiting room during the experiment. The child was given an age-appropriate description of the experiment and asked for assent. At the end of the study, each child compensated with a £15 gift voucher and a factsheet about the brain to thank them for their time and effort.

## Materials

As described in Elward & Vargha-Khadem (2018), four videos were created that contain semantic information on a particular topic. Each video contained narration that was comparable on word length (189–190 words) and reading level (Flesch Reading Ease score ranged from 60.9–64.2). A complete transcript of each of the videos and the 20 memory test questions are provided in supplementary materials. This text was animated with cartoon drawings to produce four videos which ranged in length from 72–84seconds. Two videos were presented under the recall-based learning condition and two were presented in the recognition-based learning condition.

### Design & Procedure

An overview of the experimental design is provided in figure 3. The experiment was designed to compare two learning conditions while controlling for order effects. To achieve this, each child attended the laboratory three times. Each visit was spaced one week apart. Participants learned two videos per session. There was a short delay memory test in the same session and a delayed memory test the following week. In the third session, children completed a neuropsychological assessment. Each appointment lasted approx. 2.5 hours.

**Figure 3:**
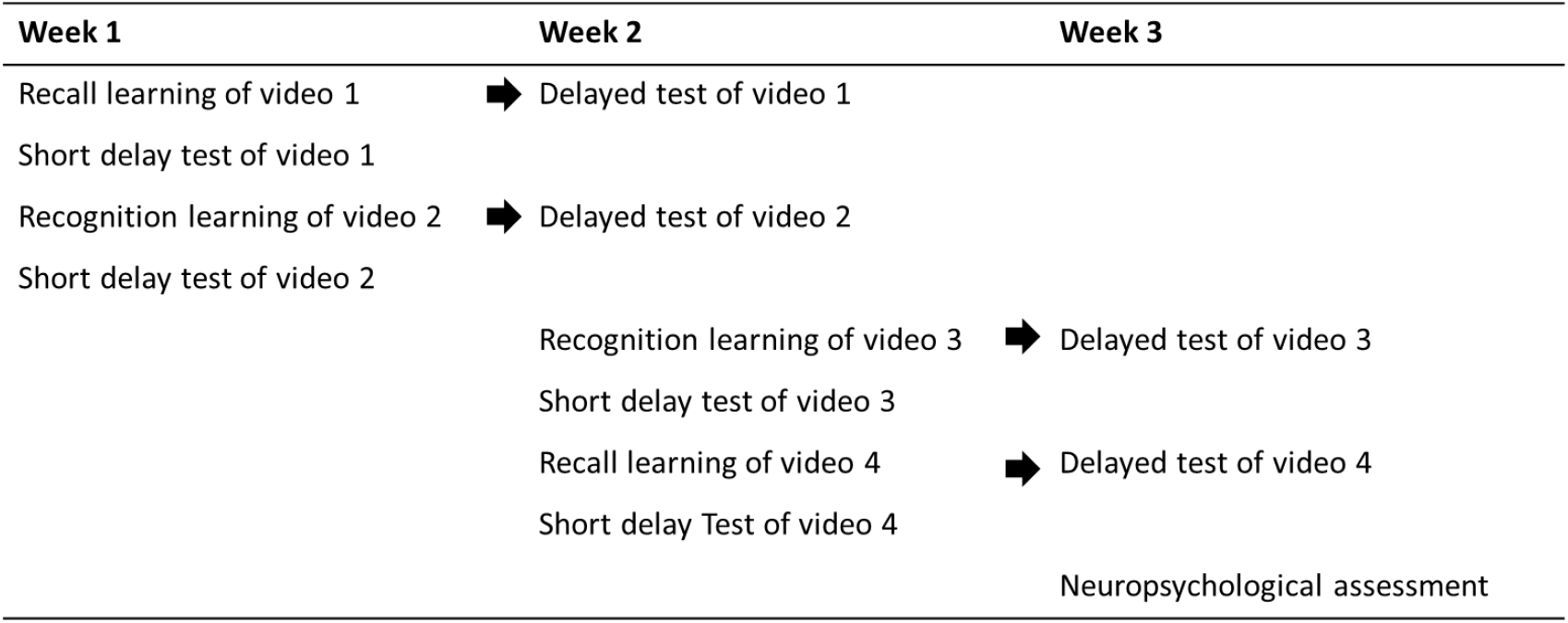
Overview of experimental design.

### Session 1

The first video was presented in the recall learning condition. A video was presented on a laptop computer. After the video, the child was presented with a recall test to facilitate learning. This test consisted of open-ended questions such as “What sort of machine was discovered in the Viking home?”. Twenty such questions were presented on PowerPoint slides. The child responded verbally, and the researcher recorded the child’s answer on a response sheet then moved the presentation to the next slide. There was no time limit for responding. The researcher encouraged the child to guess if they weren’t sure of the answer. After the test was complete the same video was played again. Six study-test cycles were completed (See figure 4: Recall-Based learning). After the six learning trials, there was a 15 minute break before the short delay test.

**Figure 4:**
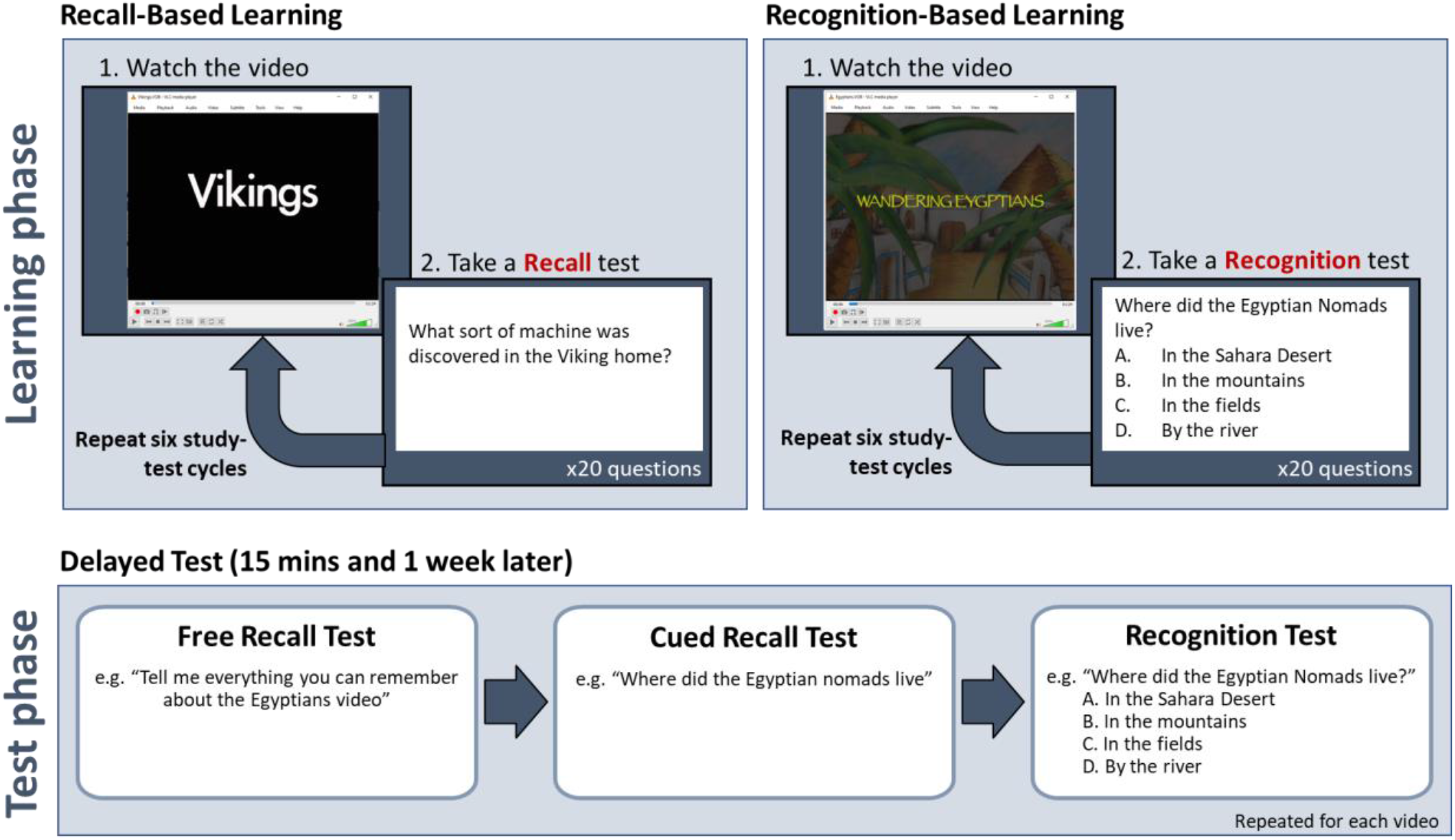
Schematic of the protocols

The short-delayed test had three stages. First, there was a free-recall test, in which the child is asked to recall anything they could remember about the video. Second, there was a cued recall test in which the child is asked the same 20 questions that they were asked in the recall learning phase. Finally, there was a recognition test where the same 20 questions are presented again but with multiple-choice response options (e.g. “What sort of machine was discovered in the Viking home? A. A Stove, B. A Loom, C. A Plough, D. A set of scales. See figure 3: Test Phase). This was followed by a self-paced break.

The second video was presented in the recognition learning condition (See figure 3: Recognition-Based learning). The recognition learning condition is identical to the recall learning condition except that after each video the child is presented with a multiple-choice test instead of a recall test. The questions were presented on PowerPoint slides. The researcher read out the questions. The child responded verbally at their own pace and was encouraged to guess if they were not sure. The researcher recorded the child’s responses. After six learning trials, there was a 15-minute break followed by a short-delay test which followed the same procedure as in the recall learning condition; first there was a free recall test, then a cued recall test and then a recognition test.

### Session 2

One week later, the child returned to the lab for the second session. At the start of this session, the child was tested on the material that they learned in the previous week. This is the 1-week delayed test and follows the same protocol as the short delay test. The test started with the free recall test, then a cued recall test and then a recognition test.

Then the child was asked to learn information contained in two more videos. The order of the learning conditions was counterbalanced so that the child learned the third video via recognition learning and the fourth video via recall learning. This is the reverse of the order in week 1 where the recall learning condition came first. Apart from the change in order, the procedure for the learning conditions and tests is identical to week 1.

To ensure that the controls undergo the same protocols as Patient H the videos are taught in the same order (Video 1 = Vikings, Video 2 = Mistletoe, Video 3 = Egyptians, Video 4 = Presidents).

### Session 3

In the third session, the child completed a 1-week delayed test for information that was learned in the second session. Lastly, control children completed a neuropsychological assessment which included the WAIS-II and the Children’s Memory Scale.

### Data Analysis

All data analysis and visualisation were conducted in R. The code and data are available on the open science framework (https://osf.io/ks3mq). Performance was averaged together to give a mean score for memory performance in the recall learning condition (videos 1 & 4) and the recognition learning condition (videos 2 & 3). The single case data are compared to controls using a Crawford-Garthwaite (2007) Bayesian test for single-case analysis (Crawford & Garthwaite, 2007) using the “psycho” package for R (Makowski, 2018).

## Results

### Learning Trials

During the learning trials, Patient H underperformed relative to controls on all recall tests but performed well on the recognition tests. The results are plotted in Figure 4.

As well as underperforming on the recall test, the patten of responses of Patient H were remarkably inconsistent. Rather than gaining knowledge incrementally though repeated presentations of the material, Patient H remembered different answers on each trial. For example, to the question “What kind of Viking home has recently been discovered?”, Patient H recalled “a farm” correctly on the first two learning trials but did not recall that detail again for the rest of the test. In response to the question, “how did the weather change when the Vikings lived in Greenland?” Patient H was alternated his response over the six learning trials so that he recalled “it got hotter” and “it got colder” three times each. However, he always correctly recalled that the animals were brought inside to protect them from the cold, indicating some memory that cold weather was an issue for the Viking settlers. Generally, the pattern of responses indicate that Patient H had a qualitatively different memory of the learning material to the typically developing children.

### 15-Minute Delayed Test

In the 15-minute delayed test, there was no clear effect of learning condition. Patient H underperformed relative to controls in the Free Recall and the Cued Recall test but performed well in the recognition memory test (see Figure 5).

**Figure 5:**
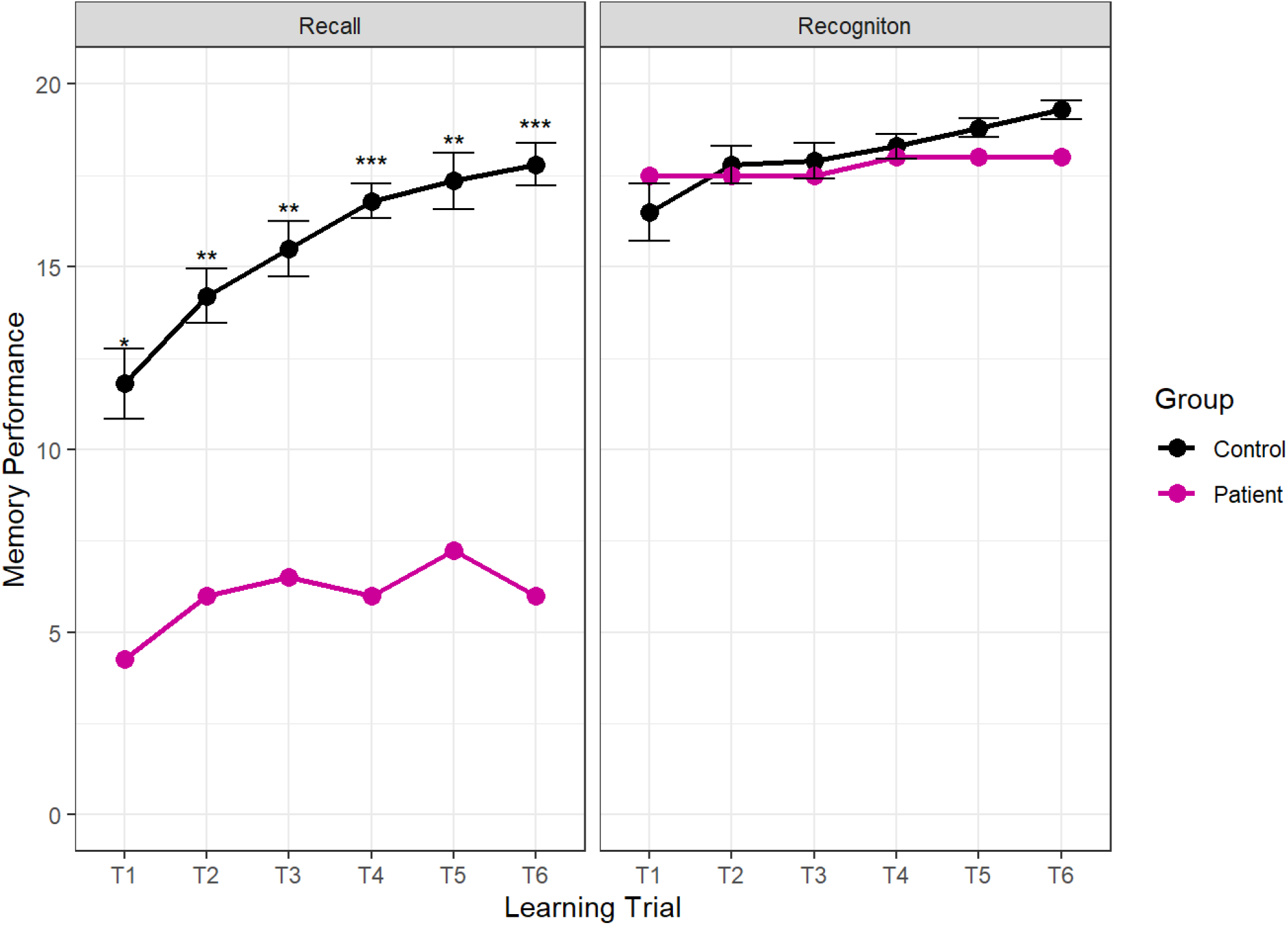
Mean memory performance across the two learning conditions. Error bars indicate 1+/ − the standard error of the mean. The outcomes of the case-control statistical tests are indicated with symbols, *p<0.05, **p<0.01, ***p<0.001

**Figure 6:**
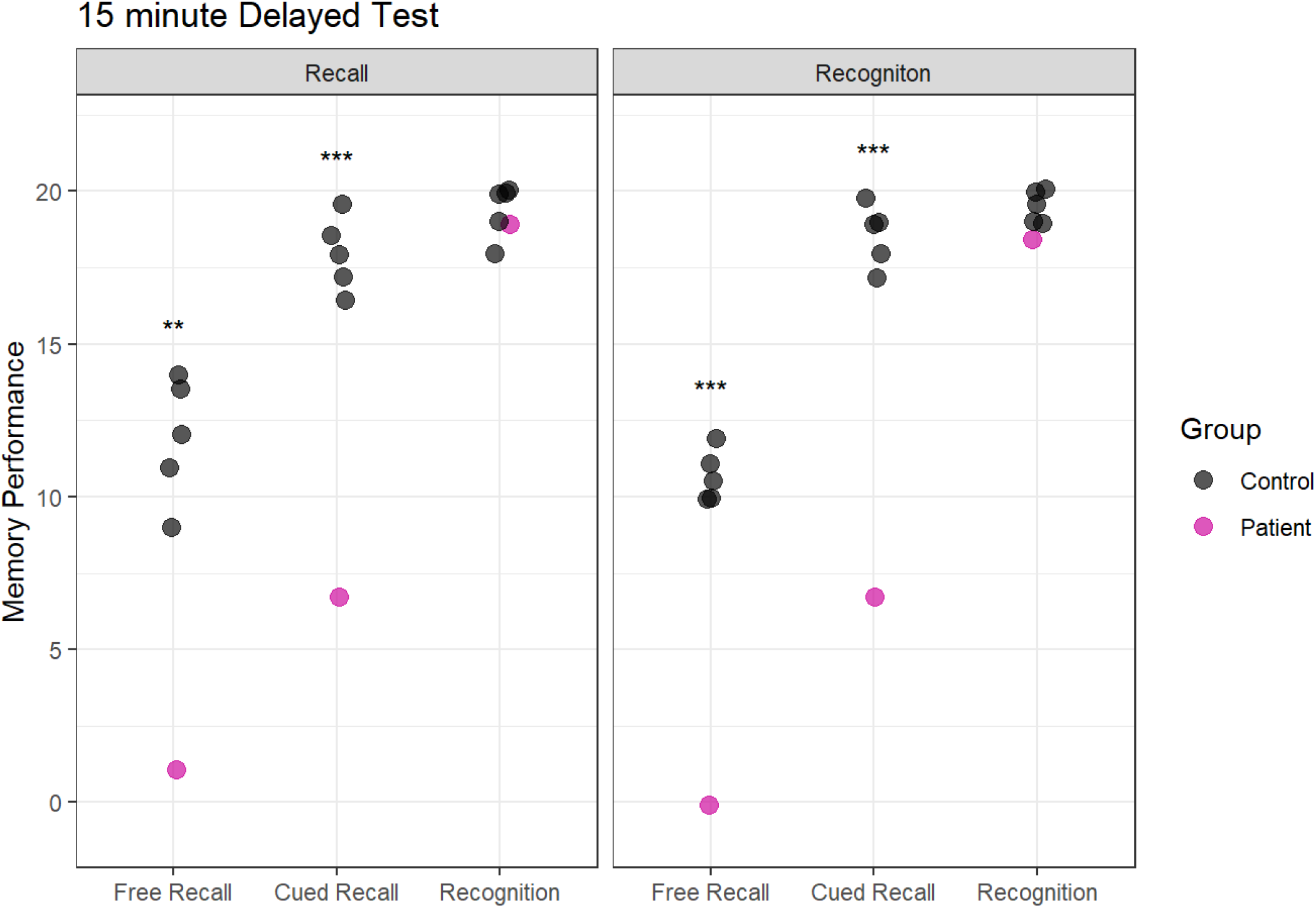
Memory performance in the 15-Minute Delayed Test following recall learning (left panel) and recognition learning (right panel). The outcomes of the case-control statistical tests are indicated with symbols, *p<0.05, **p<0.01, ***p<0.001

**Figure 7:**
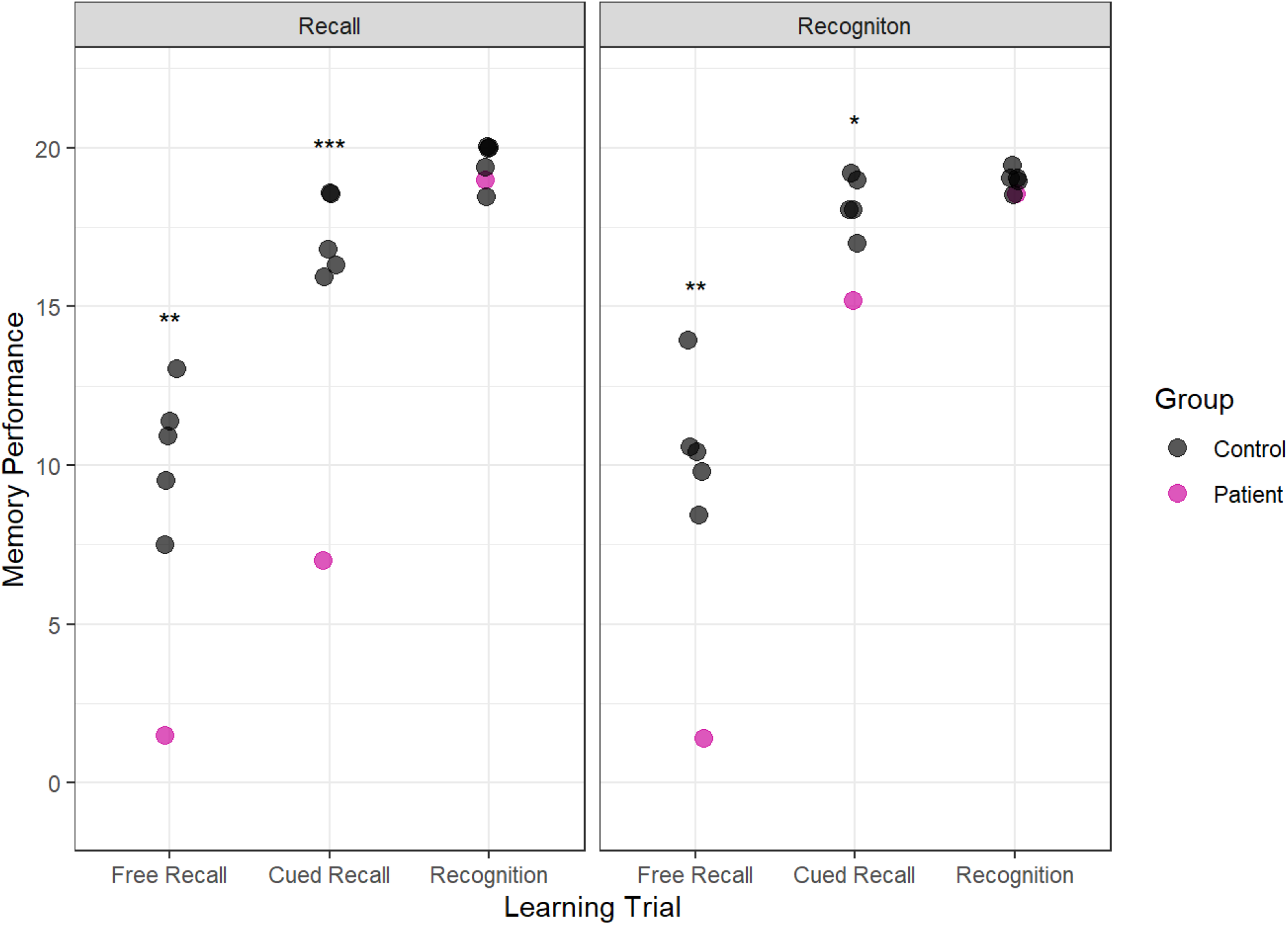
Memory performance in the 15-Minute Delayed Test. The outcomes of the case-control statistical tests are indicated with symbols, *p<0.05, **p<0.01, ***p<0.001

### 1-week Delayed Test

The crucial test was conducted one week later when we assessed whether recall-based learning or recognition-based learning was associated with better performance after a long delay. The results show that Patient H was able to recall more than twice as much information that was presented via recognition-based learning (15.25) than recall-based learning (7.00). Importantly, this was true when the learning was assessed with a cued recall test, indicating that recognition during learning facilitated the formation of a memory that could be recalled after a delay.

To compare learning across the two conditions, a difference score was computed between Patient H’s score in the cued recall test and each of the controls. In the recall testing condition, Patient H scored an average of 10.2 points lower than the control group (Standard Deviation = 1.2) but in the recognition testing condition, Patient H scored only 3 points below the control group (Standard Deviation = 0.90). A within-subjects t-test confirmed that the difference between Patient H’s data and the controls data was significantly reduced in the recognition learning condition compared to the recall learning condition t (4) = 12.24, p < 0.001.

## Discussion

These results build on the findings from Limond and colleagues that recognition testing can support semantic learning in DA (Elward & Vargha-Khadem, 2018). The previous studies showed impressive learning in adults with DA following repeated recognition testing, however, the interpretation of the previous findings were limited by changes in methodology across the experiments. Here, the methodology was controlled to allow a direct comparison of recognition testing and recall testing. The paradigm was employed with a patient and controls who had not been exposed to this material before. Although this is a single case of DA, it provides evidence that recognition testing can support semantic learning in children with DA.

In the introduction, we posited that the key to supporting semantic learning in DA may be to consider how a new semantic memory can be strengthened so that it is available for delayed recall. One mechanism by which new memories are strengthened and stabilised into semantic memory is via reconsolidation (Dudai, 2004, 2012; Squire et al., 2015). In typically developing people, new declarative memories are thought to be labile and vulnerable to forgetting. Reconsolidation occurs when the memory trace is reactivated (or replayed during sleep). Each reactivation of the prior memory is thought to create a new memory representation in the cortex which overlaps with the original memory. This process enables commonalities across events to be extrapolated and an acontextual semantic memory can form. This acontextual memory is more stable and resistant to forgetting than the episodic memory for the individual learning events.

Reactivation and consolidation are thought to be crucially dependent on the hippocampal complex which is severely compromised in patients with DA. However, recent research from our laboratory indicates that patients with DA can reactivate prior events in memory even when that information is unavailable to recall (Elward et al., 2021). This is consistent with several other accounts that have demonstrated that hippocampal processes may not be strictly necessary for reinstatement of a prior event in memory (Gagnon et al., 2018; Thakral et al., 2017). If hippocampal processing is not crucial for reinstatement of a prior learning event, patients with DA may make use of reconsolidation to strengthen semantic memories without conscious memory of the previous event. Further research combining reinstatement and semantic learning in patients with DA may test this hypothesis.

The phenomenon that retrieval practice supports learning has been referred to as ‘the testing effect’ (Roediger & Butler, 2011). In typically-developing people, active and elaborative re-processing of newly learned information, such as taking a revision test, is associated with better performance than restudying the information. This is because the process of memory retrieval is thought to support reconsolidation. However, the type of retrieval test is important for testing to facilitate learning. In research with typically-developing young adults, taking a multiple choice revision test did not support learning any more than restudying the information (Kang et al., 2007). The benefit of testing only became apparent with more elaborative tests, including short answer questions. Kang and colleagues interpret that the more demanding the retrieval process, the greater the benefit to learning. Unfortunately, elaborative tests and short answer questions are too demanding for patients with DA and are ineffective for learning in this patient group. Short answer questions require recall and patients with DA have particular difficulty with recall which is dependent on the integrity of the hippocampus (Patai et al., 2015). Patients with DA, however, have a remarkably preserved ability to recognise familiar items (Adlam et al., 2009). The data presented here indicate that recognition may be used to support semantic consolidation in this patient group. Moreover, our findings can be interpreted with in the hierarchical model of memory whereby episodic memory and recall are crucially dependent on the hippocampus, but recognition memory and semantic memory are supported by cortical structures (Mishkin et al., 1997). Learning in DA is optimal when patients can utilise cortical processing of the learned information (i.e. recognition) and avoid processing the information via the episodic memory system (i.e. recall) which is damaged in patients with DA.

This report is not the first to demonstrate that semantic processing of studied material benefits learning in DA. In a previous study with patient Jon, an adult patient with DA, a more elaborative, semantic encoding strategy (rating the words for their pleasantness) was associated with better memory performance than perceptual processing (counting the number of syllables in the words) (Gardiner et al., 2006). Therefore, patients with DA may benefit from semantic processing of the studied material in multiple choice tests. This is more demanding than passive re-exposure to the studied materials, but not so demanding that it is impossible for the patients to succeed with the test (viz recall testing, or short answer questions).

We also reported some qualitative observations of learning in our patient with DA. We described that he did not learn information incrementally such that he maintained some information that he had learned on previous trials and added additional information with more learning experiences. Instead, we report a highly variable set of responses where some information was forgotten and other information was recalled on each recall learning trial. A similar patten can be seen in the CAVLT-2 (see Table 1: Neuropsychological Assessment) where Patient H fails to learn new items on each learning trial but consistently recalls around 5 items each time. This pattern is not seen in typically developing people. Gardiner et al., (2008) described this phenomenon as ‘intertrial-forgetting’ and demonstrated that controls rarely forgot items between learning trials, but this was significantly more common the DA patient Jon. Therefore, although semantic memory is a strength in people with DA, the learning of new semantic information is inconsistent and piecemeal. Typically developing people can use the episodic memory system to scaffold learning. Children with DA experience learning in a qualitatively different way to typically developing people and must be supported differently in order to succeed in education.

**Table 1:** Neuropsychological Assessment Results of Patient H

| *Test* | *Standard Score* | *Percentile* |
| --- | --- | --- |
| **Wechsler Intelligence Scale for Children (IV)** |  |  |
| Full Scale IQ | 116 | 86 |
| Verbal Comprehension | 114 | 82 |
| Perceptual Reasoning | 121 | 92 |
| Working Memory | 113 | 81 |
| Processing Speed | 94 | 34 |
| **Children's Memory Scale** |  |  |
| Visual Immediate | 115 | 84 |
| Visual Delayed | 109 | 73 |
| Verbal Immediate | 66 | 1 |
| Verbal Delayed | 54 | 0.1 |
| General Memory | 80 | 9 |
| Attention / Concentration | 115 | 84 |
| Learning | 78 | 7 |
| Delayed Recognition | 82 | 12 |
| **One-Word Picture Vocabulary Test** |  |  |
| Expressive | 124 | 95 |
| Receptive | 99 | 49 |
| **Children's Auditory Verbal Learning Test (CAVLT-2)** |  |  |
| Immediate Memory | 89 | 23 |
| Level of Learning | 66 | 1 |
| Interference | 83 | 13 |
| Immediate Recall | < 60 | < 1 |
| Delayed Recall | < 60 | < 1 |
| Learning Trial 1 | 100 | 50 |
| Learning Trial 2 | 85 | 16 |
| Learning Trial 3 | 67 | 1 |
| Learning Trial 4 | 76 | 5 |
| Learning Trial 5 | 65 | 1 |
| **Doors and People** |  |  |
|  | **Raw Score** | **Percentile** |
| People (Recall) | 16/36 | 25 |
| Doors (Recognition) | 16/24 | 50 |
| Shapes (Recall) | 16/36 | 25 |
| Names (Recognition) | 20/24 | 95 |

Irrespective of the mechanism that supports semantic learning, the data reported here indicate that multiple choice tests can have a large effect on learning outcomes in children with DA. Multiple choice tests and pop quizzes can be readily incorporated into the classroom environment and are widely available in revision guides and educational websites. Once an appropriate test has been identified, the child may take the same test multiple times and this may improve the child’s ability to keep up with peers and remain in education to their full potential, despite their injury. Educational professionals who are supporting a child with amnesia could make this reasonable adjustment to homework or classroom quizzes so that children with DA can participate more fully in education.

## Supporting information

Supplemental Materials Video Transcript

## Notes

### Competing Interest Statement

The authors have declared no competing interest.

