## Supplemental Materials Video Transcript for "Using Recognition Testing to Support Semantic Learning in Developmental Amnesia"

### **Supplementary Materials**

### Video 1: Vikings

Running time: 00:01:24

Word count: 190

Flesch Reading Ease score: 63.3

##### Transcript

Archaeologists have recently discovered a Viking farm buried under the sand in Greenland. They have found pieces of a loom used for weaving cloth and other household belongings, including an iron knife, stone pots and a comb. The Vikings also left behind arrows made from iron and antlers weapons needed for protecting themselves and their cattle from wild animals. When the Vikings first arrived in Greenland, they did well at farming and traded with the sailors, who passed by. They sold walrus ivory and animal skins to buy timber and iron. Over the years, the sailors stopped traveling to Greenland and the Vikings could no longer make the tools that they needed for farming and building. The weather in Greenland slowly changed. The summers became shorter and colder. The short summers made it hard for the farmers to grow their crops and provide enough winter food for the cattle. As it got colder, the Vikings altered their houses from their original large single room, they created several small rooms that were warmer. The Viking farmers also moved their cattle indoors so that they would benefit from the animals’ body heat.

Video 2: Mistletoe

Running time: 00:01:12

Word count: 189

Flesch Reading Ease score: 64.2

##### Transcript

Using shotguns to harvest mistletoe is an age-old tradition in the united states. People use this peculiar method because they can't reach the mistletoe any other way, because it grows by attaching itself to the top branches of mature trees. Shooting mistletoe is good for trees as mistletoe can kill them if it grows too much. Mistletoe hunters have to be careful not to damage the tree or the mistletoe when they shoot it. They're not just trying to save the trees, but want the mistletoe in large clumps that they can then sell on to flower shops for Christmas decorations. Mistletoe has always been popular in America. The ancient Druids considered mistletoe to be sacred, because when other plants turn brown and drop their leaves, mistletoe stays green. The Druids believed that could cure illness, counteract poisons and witchcraft. When enemies happened to meet under mistletoe in the forest, they had to lay down their weapons and observe a truce until the next day. In America, during difficult times, it became a symbol of strength as it stubbornly thrived whilst most plants were dying from lack of water.

### Video 3: Egyptians

Running time: 00:01:24

Word count: 190

Flesch Reading Ease score: 63.8

##### Transcript

The ancient Egyptians are well known for their spectacular cities and pyramids. Surprisingly, thousands of years before they built these architectural masterpieces, the Egyptian people were nomads who roamed in the harsh and arid climate of the Sahara desert. In the rocky part of the desert, they left behind pictographs and rock paintings, proving that they once lived there. Today it is difficult for plants and animals to survive in the sandy part of the desert, but the nomadic Egyptians did live there. They left behind stone tools that archaeologists recognize as blades used to cut and harvest grass. In the time when the Egyptian nomads lived in the desert, it was wetter than it is now. It was still very hot and dry, but occasional monsoon rains covered the sand with pools of water. This small amount of rainwater was enough for grass to grow. Even today, the landscape is made of sand ripples rather than the more familiar dunes. Each ripple is 1 to 2 meters high and between the ripples are depressions where the rainwater must have pulled and hence where grass seeds would collect and grow.

### Video 4: Presidents

Running time: 00:01:19

Flesch Reading Ease score: 60.9

##### Transcript

In Great Britain, ex-prime ministers often return to their previous jobs in politics. They will return to the House of Commons and carry on working for their political party. In America, ex-presidents will usually leave politics to do other things. Former presidents often retire to a quieter life, writing books about what they did as president. Things were not so easy 200 years ago when it was an act of selfless public service to be president. Presidents were not paid, and many came close to bankruptcy, which is what happened to Thomas Jefferson who left the presidency heavily in debt. He had to sell his whole book collection to pay off what he owed, and these books were used to start a famous library. He still wanted to help the American people, and so he decided to help set up a university. He designed and supervised the construction of the university buildings. He set the courses and chose the staff to work there. His interest in education and learning continued for many years, but never allowed him to earn enough money to live without being in debt.
